## Supplemental Tables 1 and 2 for "High-resolution secretory timeline from vesicle formation at the Golgi to fusion at the plasma membrane in S. cerevisiae"

Table 1 - Yeast Strains - Gingras et al. 2022

| Yeast ID | Brief Description | Mating | Source | Abbreviated Genotype |
|---|---|---|---|---|
| **BY4741** | Wildtype a **(Backgrounds of all others)** | a | Brachmann et al., 1998 | *his3Δ1 leu2Δ0 ura3Δ0 met15Δ0* |
| **BY4742** | Wildtype α **(Backgrounds of all others)** | α | Brachmann et al., 1998 | *his3Δ1 leu2Δ0 ura3Δ0 lys2Δ0* |
| **9274** | mNG-Ypt31/mNG-Ypt31 | Diploid | This Study | *mNG-Ypt31::Leu2/mNG-Ypt31::Leu2* |
| **9559** | mScarlet-Ypt31/mScarlet-Ypt31 | Diploid | This Study | *mScarlet-Ypt31::Leu2/mScarlet-Ypt31::Leu2* |
| **9547** | mScarlet-Ypt31/mScarlet-Ypt31 GFP-Sec4/Sec4 | Diploid | This Study | *mScarlet-Ypt31::Leu2/mScarlet-Ypt31::Leu2 GFP-Sec4::Ura/Sec4* |
| **9568** | mScarlet-Ypt31/mScarlet-Ypt31 mNG-mCC-Myo2 Marker | Diploid | This Study | *mScarlet-Ypt31::Leu2/mScarlet-Ypt31::Leu2 mNG-mCC-Myo2(CBD)::His3* |
| **9569** | mScarlet-Ypt31/mScarlet-Ypt31 Sec2-mNG/Sec2-mNG | Diploid | This Study | *mScarlet-Ypt31::Leu2/mScarlet-Ypt31::Leu2 Sec2-mNG::Ura3/Sec2-mNG::Ura3* |
| **9548** | mScarlet-Ypt31/mScarlet-Ypt31 mNG-Ypt32/mNG-Ypt32 | Diploid | This Study | *mScarlet-Ypt31::Leu/mScarlet-Ypt31::Leu mNG-Ypt32::Leu/mNG-Ypt32::Leu* |
| **9172** | Sec7-mNG | a | This Study | *Sec7-mNG::Leu2* |
| **9173** | Sec7-mNG | α | This Study | *Sec7-mNG::Leu2* |
| **XXXX** | Sec7-mNG/Sec7-mNG | Diploid | This Study | Unstored mating of 9172 and 9173 (above) |
| **9497** | mScarlet-Ypt32/mScarlet-Ypt32 GFP-Sec4/Sec4 | Diploid | This Study | *pYpt1-mScarlet-Ypt32::Leu2/pYpt1-mScarlet-Ypt32::Leu2 GFP-Sec4::Ura3/Sec4* |
| **9498** | mScarlet-Ypt32/mScarlet-Ypt32 Sec2-mNG/Sec2-mNG | Diploid | This Study | *pYpt1-mScarlet-Ypt32::Leu2/pYpt1-mScarlet-Ypt32::Leu2 Sec2-mNG::Ura3/Sec2-mNG::Ura3* |
| **9561** | mNG-mCC-Myo2 Marker | a | This Study | *His3::mNG-mCC-Myo2(CBD)* |
| **9195** | Sec3-mNG | a | This Study | *Sec3-mNG::Ura3* |
| **9454** | Sec3-mNG/Sec3-mNG | Diploid | This Study | *Sec3-mNG::Ura3/Sec3-mNG::Ura3* |
| **9039** | Sec3-GFP/Sec3 | Diploid | This Study | *Sec3-GFP::His/Sec3* |
| **9040** | Sec5-GFP/Sec5 | Diploid | This Study | *Sec5-GFP::His/Sec5* |
| **9281** | Sec5-mNG | a | This Study | *Sec5-mNG::Ura3* |
| **9288** | Sec6-mNG | a | This Study | *Sec6-mNG::Ura3* |
| **9289** | Sec8-mNG | a | This Study | *Sec8-mNG::Ura3* |
| **9290** | Sec10-mNG | a | This Study | *Sec10-mNG::Ura3* |
| **9282** | Sec15-mNG | a | This Study | *Sec15-mNG::Ura3* |
| **9291** | Exo70-mNG | a | This Study | *Exo70-mNG::Ura3* |
| **9283** | Exo84-mNG | a | This Study | *Exo84-mNG::Ura3* |
| **9294** | Sec15-mNG | α | This Study | *Sec15-mNG::Ura3* |
| **4660** | Exocyst-3x-mNG | α | This Study | *Exo84-mNG::Ura3 Sec15-mNG::Ura3 Sec10-mNG::Ura3* |
| **4662** | Exocyst-3x-mNG | a | This Study | *Exo84-mNG::Ura3 Sec15-mNG::Ura3 Sec10-mNG::Ura3* |
| **XXXX** | Exocyst-3x-mNG | Diploid | This Study | Unstored mating of 4660 and 4662 (above) |
| **9557** | Exocyst-3x-mScarlet (mS) | a | This Study | *Exo84-mS::NatMX Sec15-mS::NatMX Sec10-mS::NatMX* |
| **9558** | Exocyst-3x-mScarlet (mS) | α | This Study | *Exo84-mS::NatMX Sec15-mS::NatMX Sec10-mS::NatMX* |
| **9565** | Exocyst-3x-mS/Exocyst-3x-mS mNG-mCC-Myo2 Marker | Diploid | This Study | *Exocyst-3x-mS::NatMX/Exocyst-3x-mS::NatMX mNG-mCC-Myo2(CBD)::His3/his3Δ* |
| **9042** | GFP-Sec4 | a | This Study | *GFP-Sec4::Ura3* |
| **3410** | GFP-Sec4 | α | Donovan and Bretscher, 2012 | *GFP-Sec4::Ura3* |
| **9043** | GFP-Sec4/Sec4 | Diploid | This Study | *GFP-Sec4::Ura3/Sec4* |
| **9044** | GFP-Sec4/GFP-Sec4 | Diploid | This Study | *GFP-Sec4::Ura3/GFP-Sec4::Ura3* |
| **9191** | GFP-Sec4 vps1Δ | α | This Study | *vps1Δ::HisMX GFP-Sec4::Ura3* |
| **3090** | Bni1-GFP/Bni1 | Diploid | | *Bni1-GFP::Ura3/Bni1* |
| **9457** | Boi2-mNG/Boi2-mNG | Diploid | This Study | *Boi2-mNG::Leu2/Boi2-mNG::Leu2* |
| **9466** | Exocyst-3x-mS/Exocyst-3x-mS Boi2-mNG/Boi2-mNG | Diploid | This Study | *Exocyst-3x-mS::NatMX/Exocyst-3x-mS::NatMX Boi2-mNG::Leu2/Boi2-mNG::Leu2* |
| **9080** | msb4Δ/msb4Δ msb3Δ/Msb3 GFP-Sec4/Sec4 | Diploid | This Study | *msb4Δ::His/msb4Δ::His msb3Δ::Leu/Msb3 GFP-Sec4::Ura/Sec4* |
| **9082** | msb3Δ/msb3Δ msb4Δ/Msb4 GFP-Sec4/Sec4 | Diploid | This Study | *msb3Δ::Kan/msb3Δ::Leu msb4Δ::His/Msb4 GFP-Sec4::Ura/Sec4* |
| **9077** | GFP-Sec4/Sec4-Q79L | Diploid | This Study | *GFP-Sec4::Ura/Sec4-Q79L::Ura* |
| **9104** | GFP-Sec4/Sec4 mCherryPM | Diploid | This Study | *GFP-Sec4::Ura/Sec4 His3::mCherryPM* |
| **9109** | msb3Δ/msb3Δ msb4Δ/Msb4 GFP-Sec4/Sec4 mCherryPM | Diploid | This Study | *msb3Δ::Kan/msb3Δ::Leu msb4Δ::His/Msb4 GFP-Sec4::Ura/Sec4 His3::mCherryPM* |
| **9118** | Gdi1/gdi1Δ GFP-Sec4/Sec4 | Diploid | This Study | *Gdi1Δ::His/Gdi1 GFP-Sec4::Ura/Sec4* |
| **9426** | Sec2-mNG/Sec2-mNG | Diploid | This Study | *Sec2-mNG::Leu2/Sec2-mNG::Leu2* |
| **9453** | Smy1-mNG/Smy1-mNG | Diploid | This Study | *Smy1-mNG::Leu2/Smy1-mNG::Leu2* |
| **9504** | Sro7-mNG/Sro7-mNG | Diploid | This Study | *Sro7-mNG::Leu2/Sro7-mNG::Leu2* |
| **9486** | Sro7-mNG/Sro7-mNG sec6-4/sec6-4 | Diploid | This Study | *Sro7-mNG::Leu2/Sro7-mNG::Leu2 sec6-4::His/Sec6-4::His* |
| **9525** | Mso1-mNG/Mso1-mNG | Diploid | This Study | *Mso1-mNG::Leu2/Mso1-mNG::Leu2* |
| **9507** | Sec1-mNG/Sec1-mNG | Diploid | This Study | *Sec1-mNG::Leu2/Sec1-mNG::Leu2* |
| **9182** | Cse4-mNG | a | Gingras et al., 2020 | *Cse4-mNG::Ura3* |
| **9315** | Rho3-imNG | α | Gingras et al., 2020 | *Rho3-imNG::Leu2* |
| **9477** | Exocyst-3x-mS/Exocyst-3x-mS GFP-Sec4/Sec4 | Diploid | This Study | *Exocyst-3x-mS::NatMX/Exocyst-3x-mS::NatMX GFP-Sec4::Ura3/Sec4* |
| **9487** | Exocyst-3x-mS/Exocyst-3x-mS Rho3-imNG/Rho3 | Diploid | This Study | *Exocyst-3x-mS::NatMX/Exocyst-3x-mS::NatMX Rho3-imNG::Leu2/Rho3* |
| **9485** | Exocyst-3x-mS/Exocyst-3x-mS Sro7-mNG/Sro7-mNG | Diploid | This Study | *Exocyst-3x-mS::NatMX/Exocyst-3x-mS::NatMX Sro7-mNG::Leu2/Sro7-mNG::Leu2* |
| **9475** | Exocyst-3x-mS/Exocyst-3x-mS Sec1-mNG/Sec1-mNG | Diploid | This Study | *Exocyst-3x-mS::NatMX/Exocyst-3x-mS::NatMX Sec1-mNG::Leu2/Sec1-mNG::Leu2* |
| **9459** | Smy1-mScarlet/Smy1-mScarlet Sec1-mNG/Sec1-mNG | Diploid | This Study | *Smy1-mScarlet::Ura3/Smy1-mScarlet::Ura3 Sec1-mNG::Leu2/Sec1-mNG::Leu2* |
| **9136** | sso2Δ | a | This Study | *sso2Δ::His* |
| **9142** | sso2Δ GFP-Sec4 | α | This Study | *sso2Δ::His GFP-Sec4::Ura* |
| **XXXX** | sso2Δ/sso2Δ GFP-Sec4/Sec4 | Diploid | This Study | Unstored mating of 9136 and 9142 (above) |
| **9099** | sro7Δ/sro7Δ GFP-Sec4/Sec4 | Diploid | This Study | *sro7Δ::KanMX/sro7Δ::KanMX GFP-Sec4::Ura3/Sec4* |
| **9121** | sec9Δ/Sec9 GFP-Sec4/Sec4 | Diploid | This Study | *Ssc9Δ::His/Sec9 GFP-Sec4::Ura3/Sec4* |
| **9141** | snc2Δ GFP-Sec4 | α | This Study | *snc2Δ::His GFP-Sec4::Ura3* |
| **XXXX** | snc2Δ/Snc2 GFP-Sec4/Sec4 | Diploid | This Study | Unstored mating of 9141 with BY4741(a) |
| **9265** | sec2Δ/Sec2 GFP-Sec4/Sec4 | Diploid | This Study | *sec2Δ::His/Sec2 GFP-Sec4::Ura3/Sec4* |
| **9122** | rho3Δ/Rho3 GFP-Sec4/Sec4 | Diploid | This Study | *rho3Δ::His/Rho3 GFP-Sec4::Ura3/Sec4* |
| **9135** | mso1Δ | a | This Study | *mso1Δ::HisMX* |
| **9137** | mso1Δ | α | This Study | *mso1Δ::HisMX* |
| **9438** | mso1Δ GFP-Sec4 | a | This Study | *mso1Δ::HisMX GFP-Sec4::Ura3* |
| **XXXX** | mso1Δ/mso1Δ GFP-Sec4/Sec4 | Diploid | This Study | Unstored mating of 9137 and 9438 (above) |
| **9117** | Sec1Δ/Sec1 GFP-Sec4/Sec4 | Diploid | This Study | *sec1Δ::His/Sec1 GFP-Sec4::Ura3/Sec4* |
| **9461** | myo2Δ/Myo2 GFP-Sec4/Sec4 | Diploid | This Study | *myo2Δ::KanMX/Myo2 GFP-Sec4::Ura3/Sec4* |
| **9456** | smy1Δ/smy1Δ GFP-Sec4/Sec4 | Diploid | This Study | *smy1Δ::KanMX/smy1Δ::His GFP-Sec4::Ura3/Sec4* |
| **9549** | msb3Δ/msb3Δ msb4Δ/Msb4 Sro7-mNG/Sro7-mNG | Diploid | This Study | *msb3Δ::Kan/msb3Δ::Leu msb4Δ::His/Msb4 Sro7-mNG::Leu2/Sro7-mNG::Ura3* |
| **9550** | msb3Δ/msb3Δ msb4Δ/Msb4 Sec1-mNG/Sec1-mNG | Diploid | This Study | *msb3Δ::Kan/msb3Δ::Leu msb4Δ::His/Msb4 Sec1-mNG::Ura3/Sec1-mNG::Ura3* |
| **9551** | msb3Δ/msb3Δ msb4Δ/Msb4 Smy1-mNG/Smy1-mNG | Diploid | This Study | *msb3Δ::Kan/msb3Δ::Leu msb4Δ::His/Msb4 Smy1-mNG::Ura3/Smy1-mNG::Ura3* |
| **9552** | msb3Δ/msb3Δ msb4Δ/Msb4 Sec2-mNG/Sec2-mNG | Diploid | This Study | *msb3Δ::Kan/msb3Δ::Leu msb4Δ::His/Msb4 Sec2-mNG::Leu2/Sec2-mNG::Ura3* |
| **9553** | msb3Δ/msb3Δ msb4Δ/Msb4 Sec15-mNG/Sec15-mNG | Diploid | This Study | *msb3Δ::Kan/msb3Δ::Leu msb4Δ::His/Msb4 Sec15-mNG::Ura3/Sec15-mNG::Ura3* |
| **9256** | mso1Δ/mso1Δ Sec1-mNG/Sec1-mNG | Diploid | This Study | *mso1Δ::His/mso1Δ::His Sec1-mNG::Leu/Sec1-mNG::Leu* |
| **9395** | Mso1(1-37)-mNG | Unknown | This Study | *Mso1(1-37)-mNG::Leu2* |
| **9556** | Sec1(1-704) | a | This Study | *Sec1(1-704)-tCYC::Ura3* |
| **9562** | mso1Δ::Mso1 | α | This Study | *mso1Δ::Mso1::His3* |
| **9563** | mso1Δ::Mso1(38-210) | α | This Study | *mso1Δ::Mso1(38-210)::His3* |

Table 2 - Plasmids Used - Gingras et al. 2022

| Plasmid | Vector | Insert | Use | First Publication |
|---|---|---|---|---|
| **4430** | pFA6a-Ura3 | *mScarlet* | Genomic tagging with mScarlet | Gingras et al., 2020 |
| **4278** | pFA6a-Leu2 | *mNG* | Genomic tagging with mNG | |
| **4279** | pFA6a-Ura3 | *mNG* | Genomic tagging with mNG | |
| **4277** | pFA6a-His3 | *mNG* | Genomic tagging with mNG | |
| **4307** | pFA6a-Ura3 | *GFP* | Genomic tagging with GFP | |
| **4354** | pFA6a-His3 | *GFP* | Genomic tagging with GFP | |
| **4600** | bRA90 | empty | Cloning of CRISPR Targets | Anand et al., 2017 |
| **4601** | bRA90 | mNG gRNA | CRISPR mNG target | This Study |
| **4611** | pFA6a-NatMX | mScarlet | mScarlet::NatMX template for CRISPR | This Study |
| **4549** | pRS315 | pYpt31-mScarlet | PCR tagging of Ypt31 | This Study |
| **4113** | pRS415 | pYpt32-mNG | PCR tagging of Ypt32 | This Study |
| **5000** | pRS415 | pYpt1-mScarlet | PCR tagging of Ypt32 | This Study |
| **5001** | pRS415 | *pCYC100-Tomato-mouseMyoVb(coiled coil)-Myo2(CBD)* | Myo2 analog | This Study |
| **5002** | pRS303 | *pCYC100-mNG-mouseMyoVb(coiled coil)-Myo2(CBD)* | His locus integrating Myo2 analog | This Study |
| **3484** | pRS415 | pTOM-mCherry-2xIST2(928-948) | PM Marker | Manford et al., 2012 |
| **4095** | pRS303 | pTDH3-mCherry-2xIst2(928-948) | His locus integrating PM marker | This Study |
| **5003** | pRS315 | *pSec4-mScarlet-Sec4(Q79L)-2xIst2(928-948)* | Constitutively-active PM-bound Sec4 | This Study |
| **16** | YEp351 | *empty* | Cloning and Control | Hill et al., 1986 |
| **4138** | YEp351 | *Sec1 Locus* | Overexpression | This Study |
| **3945** | YEp351 | *Rgd1 Locus* | Overexpression | This Study |
| **3956** | YEp351 | *Sec2 Locus* | Overexpression | This Study |
| **3857** | YEp351 | *Smy1 Locus* | Overexpression | Lwin et al., 2016 |
| **3928** | YEp352 | *Cdc42 Locus* | Overexpression | Gingras et al., 2020 |
| **3863** | YEp351 | *Rgd3 Locus* | Overexpression | Gingras et al., 2020 |
| **3854** | YEp351 | *Ypt31 Locus* | Overexpression | Gingras et al., 2020 |
| **4132** | YEp352 | *pMyo2-Myo2-3xFLAG* | Overexpression | This Study |
| **3052** | pRS425 | *empty* | Cloning and Control | Christianson et al., 1992 |
| **4432** | pRS425 | *Rho3 Locus* | Overexpression | Gingras et al., 2020 |
| **4140** | pRS425 | *Sro7 Locus* | Overexpression | This Study |
| **4153** | pRS425 | *Sec9 Locus* | Overexpression | This Study |
| **4154** | pRS425 | *Mso1 Locus* | Overexpression | This Study |
| **4447** | pRS425 | *Boi2 Locus* | Overexpression | This Study |
| **4371** | pRS425 | *Sso2 Locus* | Overexpression | This Study |
| **5004** | pRS415 | *pCYC1-mNG-Sec1(670-724)* | Sec1 C-terminus localization | This Study |
| **5005** | pRS303 | *pMso1-Mso1(38-210)* | Integration of N-terminally truncated Mso1 | This Study |
